# Sequencing and Imputation in GWAS: Cost-Effective Strategies to Increase Power and Genomic Coverage Across Diverse Populations

**DOI:** 10.1101/548321

**Authors:** Corbin Quick, Pramod Anugu, Solomon Musani, Scott T. Weiss, Esteban G. Burchard, Marquitta J. White, Kevin L. Keys, NHLBI Trans-Omics for Precision Medicine (TOPMed) Consortium, Francesco Cucca, Carlo Sidore, Michael Boehnke, Christian Fuchsberger

## Abstract

A key aim for current genome-wide association studies (GWAS) is to interrogate the full spectrum of genetic variation underlying human traits, including rare variants, across populations. Deep whole-genome sequencing is the gold standard to capture the full spectrum of genetic variation, but remains prohibitively expensive for large samples. Array genotyping interrogates a sparser set of variants, which can be used as a scaffold for genotype imputation to capture variation across a wider set of variants. However, imputation coverage and accuracy depend crucially on the reference panel size and genetic distance from the target population.

Here, we consider a strategy in which a subset of study participants is sequenced and the rest array-genotyped and imputed using a reference panel that comprises the sequenced study participants and individuals from an external reference panel. We systematically assess how imputation quality and statistical power for association depend on the number of individuals sequenced and included in the reference panel for two admixed populations (African and Latino Americans) and two European population isolates (Sardinians and Finns). We develop a framework to identify powerful and cost-effective GWAS designs in these populations given current sequencing and array genotyping costs. For populations that are well-represented in current reference panels, we find that array genotyping alone is cost-effective and well-powered to detect both common- and rare-variant associations. For poorly represented populations, we find that sequencing a subset of study participants to improve imputation is often more cost-effective than array genotyping alone, and can substantially increase genomic coverage and power.

## INTRODUCTION

Genome-wide association studies (GWAS) have detected thousands of common genetic variants associated with hundreds of complex diseases and traits^1^. A key aim for the next wave of GWAS is to interrogate the full spectrum of genetic variation underlying human genetic traits, including rare (minor allele frequency [MAF] < 0.5%) variants, across a wide range of human populations. Detecting association at rare variants requires both more comprehensive genomic coverage and sufficient sample size. Deep whole genome sequencing (WGS) is the gold standard method for capturing rare variation; however, even in the era of the $1,000 genome, large WGS association studies remain prohibitively expensive. Genotype imputation has been a mainstay of GWAS, providing increased genomic coverage from inexpensive array-based genotype call sets. While initial imputation studies only surveyed common variants (e.g.^2^), larger and more diverse reference panels now enable more accurate and comprehensive imputation of rare and low-frequency (0.5% < MAF < 5%) variants across a wide range of populations (e.g.^3^).

Imputation algorithms model haplotypes in the study sample as mosaics of haplotypes in a reference panel (e.g. from the International HapMap Project^4^ or 1000 Genomes Project^5^) to predict genotypes at untyped variants^6^. By increasing genomic coverage and accuracy, imputation increases statistical power to detect association, enables more complete meta-analysis of results from multiple studies, and facilitates the identification of causal variants through genetic fine-mapping^6; 7^. Imputation coverage and accuracy depend crucially on the size of the reference panel and the genetic distance between reference and target populations^6; 8^. The largest current broadly available reference panels, e.g. from the Haplotype Reference Consortium^9^ (HRC) and UK10K project^10^, include tens of thousands of predominantly European individuals. These panels provide near complete imputation of genetic variation down to MAF∼0.1% for many European populations, but lower imputation quality for non-European and admixed populations and population isolates, particularly for rare and low-frequency variants^11-13^. The 1000 Genomes Project and HapMap panels include individuals from diverse worldwide populations, but provide more limited imputation coverage and accuracy due to their smaller sample sizes.

Capturing rare variation across diverse populations is crucial to detect population differences in genetic risk factors, accurately predict genetic risk, and identify causal variants and biological mechanisms through trans-ethnic fine-mapping^14; 15^. Population-matched or multiethnic reference panels can improve imputation quality and coverage for rare variants in GWAS of diverse populations^11-13; 16-18^; this approach has enabled discovery of novel loci and refinement of association signals for multiple populations and complex traits^12; 19; 20^.

Here, we consider an approach in which a subset of study participants is whole genome sequenced and the rest are array-genotyped and imputed using an augmented reference panel that comprises the sequenced participants and individuals from an external reference panel^21; 22^. This hybrid sequencing-and-imputation strategy provides more comprehensive coverage than array genotyping alone, and is less costly than whole genome sequencing the entire sample. We and others have used this strategy^18; 23-25^, but no analysis of coverage, power, and cost-effectiveness has been carried out to date. Here, we assess how imputation coverage and power to detect association vary across genotyping arrays and as a functions of the number of population-matched individuals sequenced and included in the reference panel for two admixed populations (African Americans and Latino Americans) and two European population isolates (Sardinians and Finns) to identify powerful and cost-effective GWAS strategies in these populations. We also describe an interactive web-based tool to assist researchers in the design and planning of their own GWAS.

## MATERIALS AND METHODS

We first describe WGS data sources used in our analysis. Next, we describe imputation strategies, and outline procedures and imputation quality metrics to compare these strategies. Finally, we present a novel method to estimate power for the sequencing-only, imputation-only, and sequencing-and-imputation strategies. For ease of presentation, we assume a dichotomous trait and a multiplicative disease model, although our findings generalize easily to continuous traits and other genetic models.

### Data Resources

We used WGS data on 11,920 individuals to assess imputation quality across reference panel configurations and genotyping arrays for admixed populations and population isolates. For our analysis of admixed populations, we used WGS data on 3,412 African Americans (participants from the Jackson Heart Study) and 2,068 Latino Americans (participants of Puerto Rican and Mexican descent from the GALA II study and Costa Rican descent from the Genetic Epidemiology of Asthma in Costa Rica and CAMP studies) in the National Heart, Lung, and Blood Institute (NHLBI) Trans-Omics for Precision Medicine (TOPMed) WGS program. For our analysis of isolated populations, we used WGS data on 2,995 Finns (participants of the GoT2D, 1KGP, SISu, and Kuusamo studies) and 3,445 Sardinians (participants of the SardiNIA study) in the HRC.

### Procedures to Evaluate Imputation Coverage and Accuracy

We considered three imputation strategies: (1) using sequenced study participants as a study-specific reference panel, (2) using an external reference panel alone (for this comparison, the HRC or HRC subset excluding individuals from the target population), and (3) using an augmented panel that comprises sequenced study participants and an external panel.

For African Americans, who are underrepresented in the current version 1.1 of the HRC, we constructed population-specific and HRC-augmented reference panels with 0 to 2,000 African Americans. For Latino Americans, we used the same approach but restricted the study-specific panel size to ≤1,500 due to the more limited available sample of sequenced Latino American individuals. For Finns and Sardinians, which are present in the HRC, we constructed augmented reference panels that comprised the 29,470 non-Finnish or 29,020 non-Sardinian individuals in the HRC together with 0 to 2,000 Finns or Sardinians from the HRC.

For each population, each imputation strategy, and each of three commonly-used genotyping arrays (Table 1), we used sequence-based genotype calls at marker variants present on the array as a scaffold for imputation using Minimac3, masking the remaining sequence-based genotype calls^7^. We then compared the imputed genotype dosages to the true (masked) genotypes to estimate (a) imputation *r*^2^, the squared Pearson correlation between true genotype and imputed dosage, and (b) imputation coverage, the proportion of variants with imputation *r*^2^> 0.3 and minor allele count (MAC) ≥ 5 (the MAC threshold used by the HRC panel^9^) in the reference panel.

**Table 1.** Genotyping Arrays used for Comparisons

| Array | No. Marker Variants | List Cost per Sample<sup>26</sup> |
| --- | --- | --- |
| Illumina Infinium Core | 307K | \$49 |
| Illumina Infinium OmniExpress | 710K | \$94 |
| Illumina Infinium Omni2.5 | 2.5M | \$172 |

### Estimating Power to Detect Association using Empirical Imputation Quality Data

When sequenced individuals are included in the reference panel, power calculations should account for the interdependence between imputation *r*^2^ and the number of participants sequenced *n*, and for the possibility that the variant is not imputable (absent in the reference panel or not imputed due to insufficient MAC, or filtered prior to association analysis due to imputation *r*^2^ falling below a given threshold). While common variant associations are likely to be captured by LD proxy SNPs even when the causal variant is not directly genotyped or imputed, rare variant associations are much less likely to be captured by proxy SNPs^27^. Here, we assume that power to detect association for variants that are not imputable is zero. This assumption affects power calculations almost exclusively for rare variants, since common variants are almost uniformly imputable with large reference panels^7; 9^.

We assume that the *n* participants who are sequenced are randomly subsampled from the overall sample of *n* + *m* study participants, and that test statistics are calculated separately for the sequenced and imputed subsamples and combined using the effective sample size weighted meta-analysis test statistic 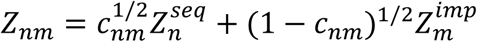, where *c*_*nm*_ = *n*/(*n* + *r*^2^*m*). The asymptotic distribution of 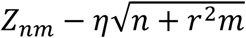 is normal with mean 0 and variance 1, where *r*^2^ is the squared correlation between imputed dosages and true genotypes, and *η* is an effect size parameter which is equal to 0 under the null hypothesis of no association. The form of *η* depends on the association model (e.g. additive, dominant, multiplicative), relative risk or odds ratio, MAF, and population prevalence and, for binary traits, the case-control ratio. Under an arbitrary association model for binary traits, we can write

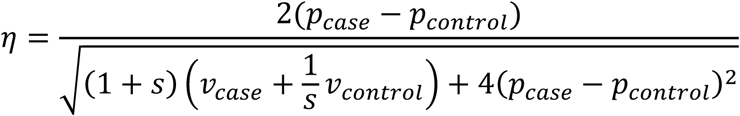

where *p*_*case*_ and *p*_*control*_ are the alternate allele frequencies in the disease-positive and disease-negative populations, *v*_*case*_ and *v*_*control*_ are the variances of genotypes in the disease-positive and disease-negative populations, and *s* is the GWAS case-control ratio.

To estimate power while accounting for variability in imputation *r*^2^ and the possibility that a variant is not imputable, we average empirical imputation *r*^2^ values and MACs across variants from experiments with real data described in the previous section. Specifically, we estimate power to detect association when *n* individuals are sequenced and *m* are genotyped and imputed as

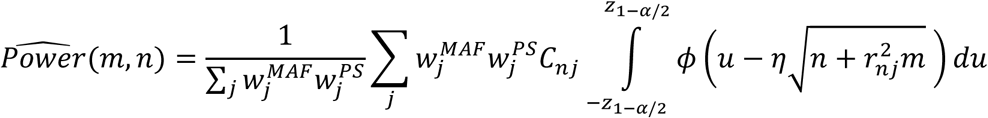

where 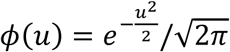 is the standard normal density function, *z*_1-*α*/2_ is the *α*-level significance threshold, 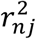 is the imputation *r*^2^ value for the *j*^*th*^ variant, 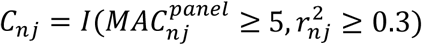 is an indicator equal to 1 if the *j*^*th*^ variant was imputable and 0 otherwise, and 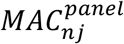 is the reference panel MAC for the *j*^*th*^ variant when the *n* sequenced individuals from the target population were included in the reference panel.

We define the first weight term 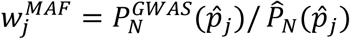, where *N* is the total number of samples used in our analysis for the given population (e.g. *N* =3,412 for African Americans),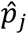 is the sample MAF for the *j*^*th*^ variant in the total sample,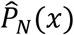 is the proportion of variants with 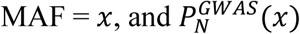 is the probability of observing sample MAF = *x* in a sample of size *N* given the specified association model. For example, in a GWAS with sample size *N* and case-control ratio *s*, the sample MAC (which is equal to 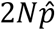, where 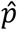 is the sample MAF) is approximately Poisson distributed with mean 2*N*(*sp*_*case*_ + *p*_*control*_)/(*s* + 1), where *p*_*case*_ = *pγ*/[1 + *p*(*γ* − 1)] and *p*_*control*_ = (*p* - *Kp*_*case*_)/(1 - *K*) for a variant with population MAF *p* and relative risk *γ* for a disease with prevalence *K*. This weighting approach adjusts for differences between the empirical distribution of MACs across variants in real data, and the theoretical MAC distribution for a variant with the specified MAF, effect size, prevalence in a GWAS with sample size *N* and case-control ratio *s*.

The second weighting term 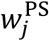 accounts for the probability that a variant with the specified population MAF *p* is population-specific (monomorphic outside the target population), and is defined

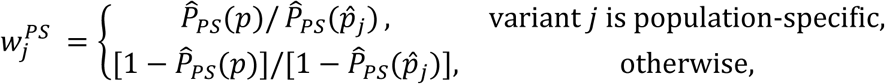

where 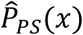 is the fraction of variants that are population-specific among variants with MAF=*x* in the target population. This adjustment factor ensures that the weight assigned to population-specific variants in power calculations reflects the probability that a variant with the specified population MAF *p* is population-specific.

## RESULTS

First, we compare strategies to improve imputation using study-specific WGS data for African Americans, Latino Americans, Sardinians, and Finns. Next, we assess the effects of genotyping array on imputation quality and coverage for each population and reference panel. We then use these results to estimate statistical power to detect association as a function of study-specific panel size, number of participants imputed, external reference panel, and genotyping array. Finally, we identify cost-effective study designs by comparing statistical power and total experimental (sequencing and genotyping) costs for sequencing-only, imputation-only, and sequencing-and-imputation GWAS designs for each population and genotyping array.

### Strategies to Improve Imputation using Study-Specific WGS Data

We compared imputation *r*^2^ and coverage (proportion of variants with imputation *r*^2^> 0.3 and reference MAC ≥ 5) for three imputation strategies: (1) using an external reference panel (the HRC or HRC subset) alone, (2) using an augmented reference panel that combines the study-specific and external panels, and (3) using a study-specific reference panel alone.

The external panel alone (HRC for Latino Americans and African Americans, and HRC subset that excludes individuals from the target population for Finns and Sardinians) provided 96% imputation coverage for MAF ≥ 0.25% variants (where MAF is calculated separately within each population) for Finns, 84% coverage for Sardinians, 86% coverage for Latino Americans, and 77% coverage for African Americans (Figure 1, top row). The relatively lower coverage for African Americans is expected since the HRC consists primarily of Central and Northern Europeans, who are genetically closer to Finns and Sardinians, and includes relatively few Africans or African Americans. Despite the small number of Latino or Native Americans included in the HRC, imputation coverage was slightly higher for Latino Americans than for Sardinians. This may reflect the high degree of European admixture in many Latino American populations^28^, and the abundance of population-specific rare and low-frequency variants in the Sardinian population^24^.

**Figure 1.**
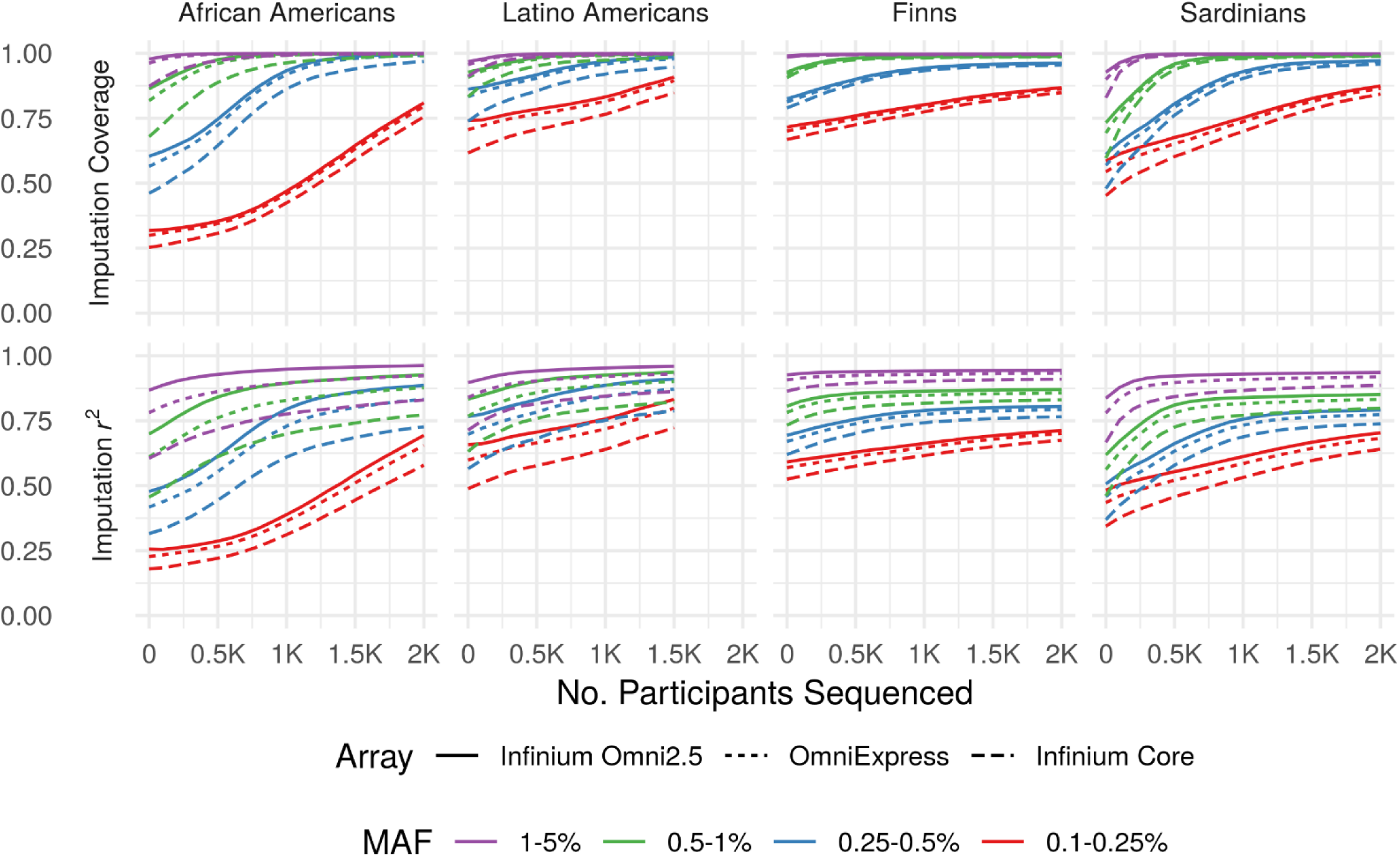
Imputation Quality by Population and Genotyping Array. Imputation coverage (upper panels) and mean imputation r^2^ (lower panels) as functions of the number of population-matched individuals included in augmented reference panels (Number Sequenced, x-axis). Here and elsewhere, MAF is calculated separately within each population.

Augmenting an external reference panel with even a relatively small number of sequenced individuals substantially increased coverage, particularly for African Americans and Sardinians, and for variants with lower MAF. For example, augmenting the external panel with 500 sequenced individuals from the study population improved overall imputation coverage for MAF=0.25-0.5% variants by 4% for Finns, 9% for Latino Americans, 16% for African Americans, and 23% for Sardinians genotyped using the OmniExpress relative to the external panel alone (Figure 1). Similarly, augmenting the external reference panel with even 200 individuals increased imputation coverage for MAF=0.1-0.25% variants by 3%, 4%, 6%, 10% relative to the external panel alone for Finns, Latino Americans, African Americans, and Sardinians, respectively.

With 2,000 individuals from the target population (or 1,500 for Latino Americans), population-specific panels provided roughly equivalent imputation *r*^2^ compared to augmented panels (Supplemental Figure 1A); however, augmented panels provided higher imputation coverage overall for low MAF variants (Supplemental Figure 1B). For example, augmented panels with 2,000 individuals from the target population (or 1,500 for Latino Americans) provided 86%, 80%, 79%, and 86% coverage for 0.1-0.25% MAF variants for Finns, Latino Americans, African Americans, and Sardinians respectively, whereas population-specific panels alone provided 72%, 51%, 78%, and 72% coverage using the Omni Express array. However, imputation coverage for variants with MAF>0.25% differed by <1% between augmented and population-specific panels with 2,000 individuals from the target population (or 1,500 for Latino Americans) for all populations and genotyping arrays. When a smaller number (less than 500) of individuals from the target population are sequenced, augmented reference panels provided substantially higher imputation coverage and *r*^2^ than population-specific panels alone. For example, augmented panels with 500 individuals from the target population provided 90%, 85%, 65%, and 85% coverage for 0.25-0.5% MAF variants for Finns, Latino Americans, African Americans, and Sardinians respectively, whereas population-specific panels of 500 individuals provided ≤ 30% coverage using the Omni Express array.

Even very rare variants (MAF=0.1-0.25%) attained high coverage across all populations given a sufficient number of population-matched individuals in the reference panel. For example, attaining ≥ 70% imputation coverage for MAF=0.1-0.25% variants required a study-specific panel of ≥ 1,800 individuals for African Americans, 1,000 for Latino Americans, 700 for Sardinians, and 0 for Finns using the OmniExpress. These increases in imputation coverage primarily reflect increasing numbers of population-specific variants captured in the reference panel, which are absent from or present in low copy number in the external panel.

### Imputation Coverage and Quality across Genotyping Arrays

Imputation coverage was generally similar for the OmniExpress and Omni2.5 arrays, but consistently lower for the less dense Core array. Coverage differed by <7% between the OmniExpress and Omni2.5 across all MAF bins, populations, and reference panels, whereas the Core provided up to 24% lower coverage than the Omni2.5 (Figure 1, upper panels). Imputation coverage was more heterogeneous across arrays for populations with greater genetic distance from the external reference panel (e.g., African Americans and the HRC panel), particularly with smaller (or absent) study-specific panels. Because we used the same reference panels for each genotyping array, differences in imputation coverage between arrays are solely due to differences in the proportion of variants that attained imputation *r*^2^ ≥ 0.3. Imputation *r*^2^ varied more across genotyping arrays than did imputation coverage (Figure 1, lower versus upper panels); however, the magnitude of differences in imputation *r*^2^ between arrays was still generally modest, particularly for the Finns and Sardinians.

### Powerful and Cost-Effective Strategies for GWAS across Populations

We compared the cost-effectiveness of sequencing-only, imputation-only, and sequencing-and-imputation strategies by analyzing statistical power to detect association as a function of numbers of study participants sequenced and imputed, genotyping array, and reference panel across a range of genetic models. Here, we define the most *cost-effective* strategy as either (1) minimizing total experimental (sequencing and genotyping) cost while attaining power at or above a given threshold, or equivalently (2) maximizing power while maintaining cost no greater than a specified constraint.

The cost-effectiveness of sequencing a subset of study participants varied greatly across populations. For Finns, imputation-only designs were most powerful to detect association and adding sequenced individuals increased power only minimally, even for low-frequency and rare variants. For Sardinians, Latino Americans, and African Americans, sequencing a subset of study participants was optimal, and often achieved substantially greater power than imputation-only or sequencing-only studies. For example, a GWAS of African Americans with equal numbers of cases and controls in which 400 participants are sequenced and 11,100 are imputed using the Illumina Infinium Core array has 90% power to detect a risk variant with MAF = 0.5% and RR = 4 for a disease with prevalence 1%, whereas an imputation-only GWAS with the same total cost (19,250 participants) has only 68% power (Figure 2). Even for populations in which optimal sequencing-and-imputation designs had substantially greater power than imputation-only, the optimal number to sequence was often modest. For example, only 210 participants are sequenced under the optimal design using the Illumina OmniExpress to attain 80% power in the previous example (Figure 3). This is expected because even a relatively small study-specific panel can substantially increase imputation coverage (Figure 1, upper panels).

**Figure 2.**
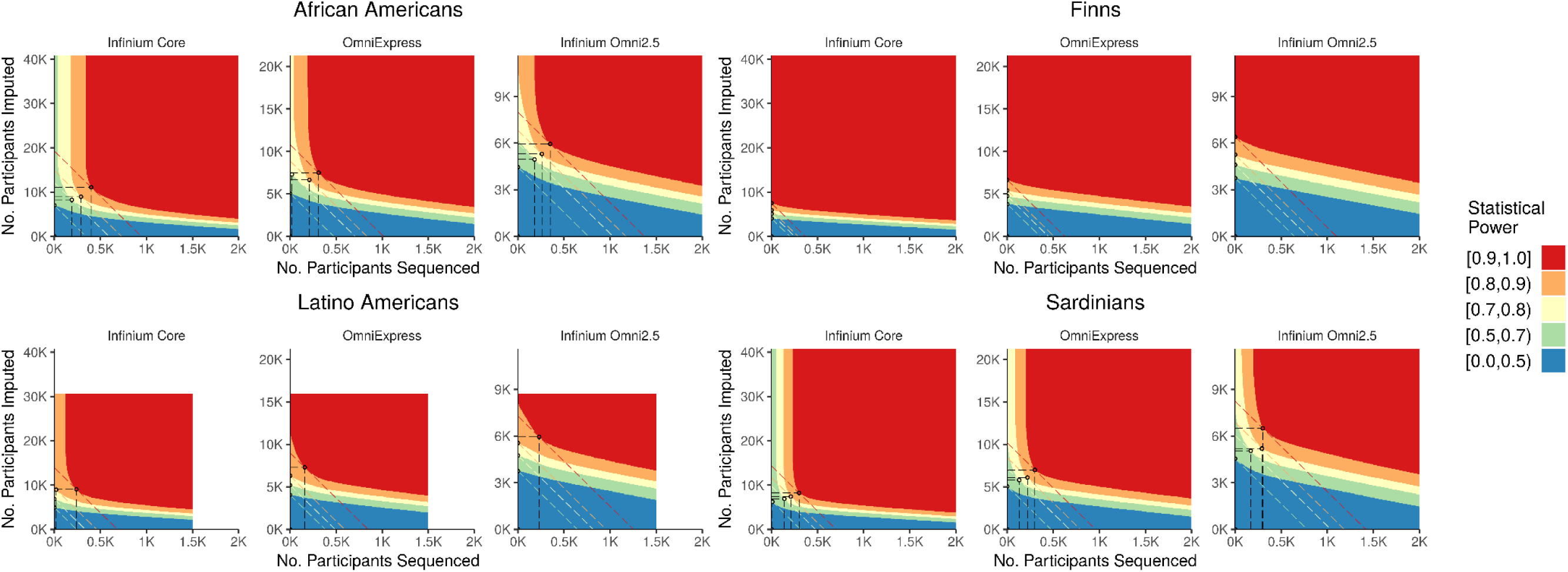
Power and Optimal Design by Population and Genotyping Array. Power to detect association for case-control studies with equal numbers of cases and controls as a function of sequenced subsample size (x-axis) and imputed subsample size (y-axis) for a variant with MAF 0.5% and relative risk 4 for a disease with prevalence 1%. Axes are scaled to reflect costs of genotyping arrays (Table 1) and sequencing ($1K per sample). Dashed diagonal lines indicate study designs with the same total cost, given by y = a - bx where a = (Total Cost) /(Array Cost) and b = (Sequencing Cost)/ (Array Cost). Circled points indicate optimal study designs, which attain the indicated power level at minimum total experimental cost (or, maximize power at the indicated total experimental cost), shown only for optimal designs with total genotyping cost ≤ $2M ($1.5M for Latino Americans).

**Figure 3.**
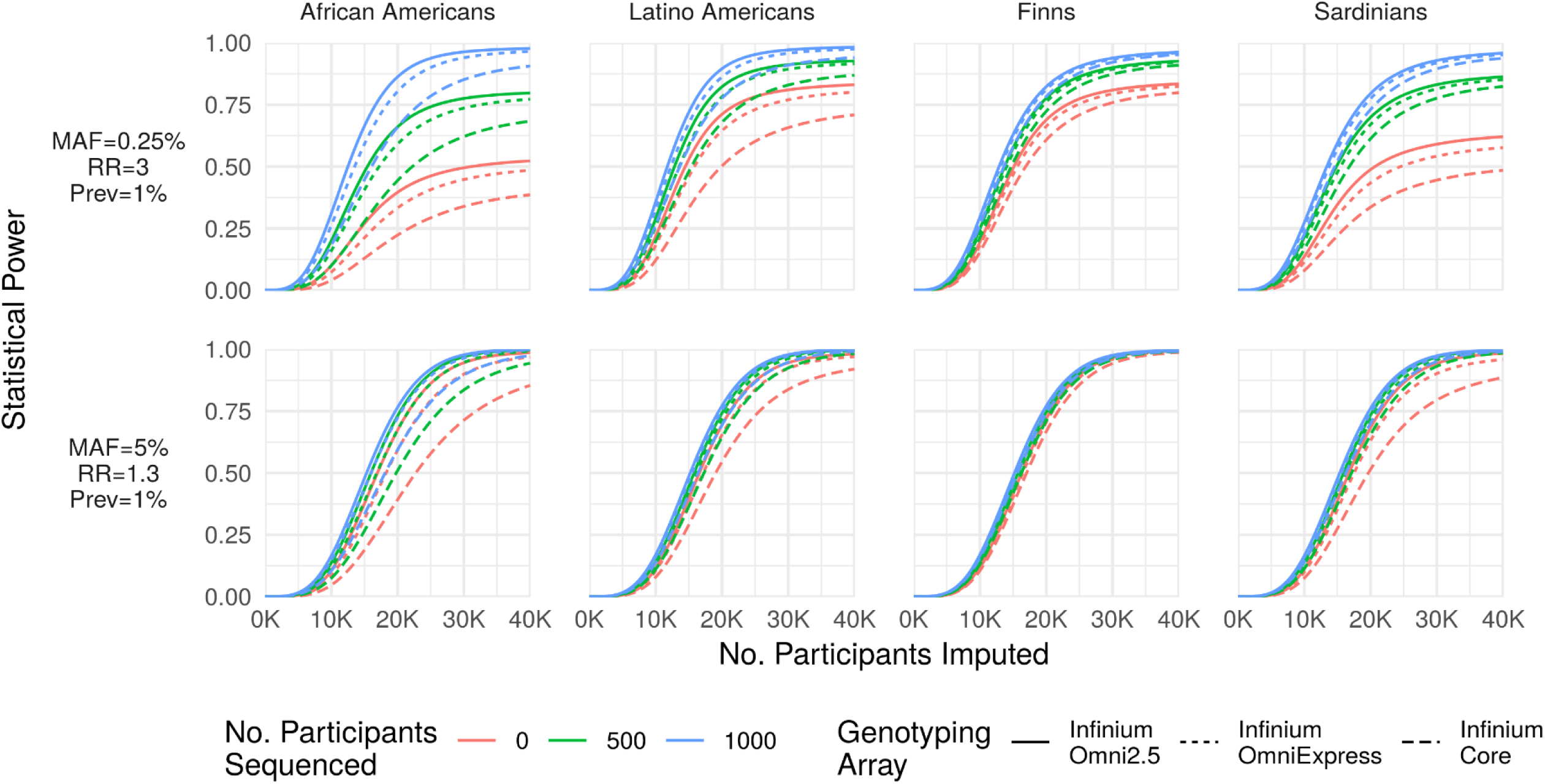
Power as a Function of Minor Allele Frequency and Effect Size. Statistical power (y-axis) to detect a rare large-effect variant (MAF=0.25%, RR=3; top row) and common modest-effect variant (MAF=5%, RR=1.3; bottom row) for a disease with prevalence 1% as a function of the number of participants array-genotyped and imputed (x-axis) when 0, 500, or 2,000 participants are sequenced and included in an augmented reference panel. The number of participants sequenced has a far greater impact on statistical power for the rare variant association. Importantly, statistical power is bounded above by the probability that the variant is imputable (r^2^ > 0.3 and reference MAC ≥ 5), causing power to asymptote below 1 as a function of the number of imputed participants (e.g., upper-left panel).

### Denser Genotyping Arrays vs. Sequencing: Which is More Cost-Effective to Increase Power?

Imputation coverage and power to detect association can be increased by using denser genotyping arrays, which provide a more informative framework for imputation, or by sequencing population-matched individuals and augmenting the reference panel. We assessed the cost-effectiveness of these two strategies by comparing power to detect association across genotyping arrays for study designs that have the same total cost assuming $1000 for WGS and current list prices for genotyping arrays (Table 1). As expected, the optimal number of participants sequenced to maximize power given fixed total cost generally decreased with increasing array density. For example, the optimal number sequenced to maximize power to detect association was 500, 300, and 90 for the Infinium Core, OmniExpress, and Omni2.5 respectively for Sardinians given total sequencing and genotyping budget of $2M for a risk variant with RR = 2, MAF = 1%, and disease prevalence 1%. Power to detect association under the optimal design given a fixed total cost was generally greater for sparser arrays; in the previous example, power under the optimal design was 98%, 91%, and 55% for the Infinium Core, OmniExpress, Omni2.5.

We also compared optimal designs to attain power above a given threshold at minimum total cost across genotyping arrays based on the per-sample array genotyping costs reported in Table 1. Generally, sparser arrays were more cost-effective (reached the power threshold with lower total cost) than dense arrays. In fact, the sparsest genotyping array in our analysis, the Infinium Core, was most cost-effective across all disease models and populations apart from African Americans, for whom the Infinium OmniExpress was most cost-effective for some rare-variant disease models. This last result is unsurprising given the substantial difference in imputation coverage between the Infinium Core and Omni arrays for African Americans (Figure 1). Importantly, our analysis assumes 1) a direct trade-off between the GWAS sample size and sequencing/array genotyping costs, and 2) no additional costs per GWAS sample other than sequencing/genotyping. Under these assumptions, we found that denser arrays are generally less cost-effective than sparser arrays; of course, denser arrays provide higher imputation coverage given a fixed GWAS sample size.

### Optimal Study Design as a Function of Minor Allele Frequency and Effect Size

Power to detect association under a given study design depends on MAF, effect size (relative risk or odds ratio), and population prevalence^29^. These parameters also influence the relative cost-effectiveness of sequencing and imputation. While common variants can be accurately imputed with small reference panels, large population-matched reference panels are needed to capture rare (population-specific) variants. In Figure 3, we illustrate the impact of sequencing on statistical power for two combinations of MAF and effect size in each of the four study populations.

The optimal percentage of study participants sequenced to attain ≥80% power to detect association at minimum total cost increases with decreasing MAF (Figure 4). This is expected, since larger reference panels are needed to capture variants with lower frequency. Finally, the optimal percentage of study participants sequenced to attain ≥80% power decreases with increasing effect size magnitude. This is expected, since the expected number of risk alleles captured in the reference panel increases with effect size magnitude.

**Figure 4.**
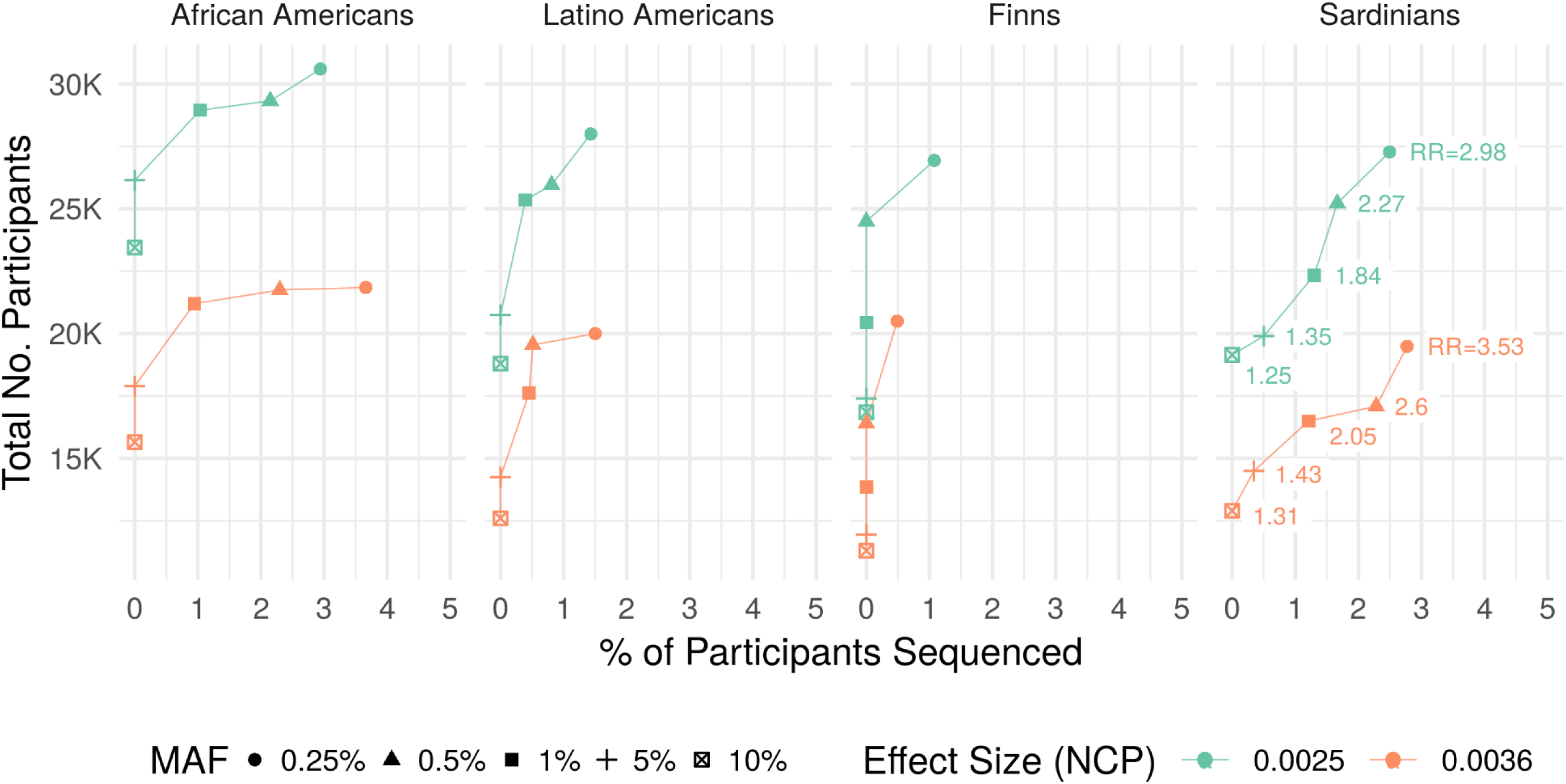
Optimal Design as a Function of Minor Allele Frequency and Effect Size. Percentage of participants sequenced (x-axis) and total sample size (y-axis) under optimal designs to attain statistical power ≥80% for rare and common variants across two effect size values for each of the four study populations using the Infinium Core array. Here, effect size refers to the χ-squared non-centrality parameter (NCP) for single-variant association tests given perfect genotype accuracy, which is defined as η^2^ in Methods. Relative risk (RR) values corresponding to each combination of MAF and NCP are indicated in the far-right panel (for Sardinians). With NCP held constant, differences in optimal design for different MAF values are solely due to differences in imputation coverage and quality across the MAF spectrum.

## DISCUSSION

While the cost of genome sequencing has fallen dramatically^29^, large genome sequencing studies remain prohibitively expensive. Large imputation reference panels are now enabling accurate imputation of even very rare variants (MAF>0.001)^9; 13; 30^, making imputation-based GWAS viable and cost-effective for detecting associations across much of the allele frequency spectrum. For populations with limited reference panel data, we have shown that sequencing a subset of study participants can substantially increase imputation coverage and accuracy, particularly for rare and population-specific variants, at a fraction of the cost of sequencing the entire study cohort. Our results also suggest that it is almost always advantageous to augment existing reference panels, except when the study-specific sequenced panel is large or the target population has high genetic distance from the external panel.

Complementary sequencing-and-imputation GWAS strategies have been applied to refine association signals and discover novel associations for several populations and complex traits^12; 19;20^. While most sequencing-and-imputation studies to date have been carried out in European isolated populations, our results suggest that this strategy can also be powerful and cost-effective for admixed and non-European populations. In addition to increasing genomic coverage and power to detect association for the study itself, sequencing a subset of study participants provides a data resource that can be used to enhance imputation in future studies of the same or related populations so long as the sequence data can be shared.

Directly augmenting an existing reference panel with study-specific sequence data is not always feasible due to technical, logistical, and privacy constraints. However, we and others have found that the distributed reference panel approach (separately imputing with two or more reference panels and combining the results) provides nearly equivalent imputation quality (Supplemental Figure 2). Thus, study-specific WGS data can be used to improve imputation even when directly augmenting an external panel is not feasible.

While large reference panels enable accurate imputation across a wide range of the allele frequency spectrum^9; 13^, the extent of genetic variation that can be captured through imputation is limited relative to WGS. For example, *de novo* mutations cannot be imputed regardless of reference panel size. This is particularly salient for monogenic disorders; for example, over 80% of achondroplasia cases occur from recurrent *de novo* mutations in *FGFR3*^31^. Thus, imputation may be unable to detect causative alleles for traits with extreme genetic architectures, even with very large reference panels.

As increasingly large and diverse sequencing projects are conducted, larger and more diverse reference panels will become available. In the design and planning of GWAS, it may be prudent to consider resources under development and pending release in addition to resources that are currently available. More broadly, our analysis highlights the utility of collaboration and coordination across institutions for effective study design and resource allocation. For example, the optimal design to maximize power in an individual study does not necessarily maximize meta-analysis power across multiple studies of the same trait and population.

Our analysis of cost-effectiveness and optimal design depends crucially on the relative per-sample costs of sequencing and array genotyping. Both sequencing and array genotyping costs have fallen markedly in recent years, and are likely to continue to do so. Depending on the relative rates of change, cost-effectiveness and optimal design also may change. In addition, the cost of participant recruitment and DNA sample collection may alter the relative cost-effectiveness of sequencing and genotyping. Finally, our cost-effectiveness analysis assumes that sample size is unconstrained; this may not apply for small populations or rare diseases.

While our results are illustrative, investigators may wish to explore questions of the relative cost-effectiveness of sequencing and array genotyping strategies in the context of their own study and relevant assumptions about population, reference panels, and sequencing and array genotyping costs. To enable this exploration, we have developed a flexible, easy-to-use tool, APSIS (Analysis of Power for Sequencing and Imputation Studies), which is open source and freely available (see Web Resources).

### Conclusions

Here, we assessed the genomic coverage, statistical power, and cost-effectiveness of sequencing and imputation-based designs for GWAS in four populations across a range of genetic models. We developed a novel method to account for available reference haplotype data in power calculations using empirical data, which can be applied to inform GWAS planning and design. For European populations that are well-represented in current reference panels, our results suggest that imputation-based GWAS is cost-effective and well-powered to detect both common- and rare-variant associations. For populations with limited representation in current reference panels, we found that sequencing a subset of study participants can substantially increase genomic coverage and power to detect association, particularly for rare and population-specific variants. Our results also suggest that larger and more diverse reference panels will be important to facilitate array-based GWAS in global populations.

## WEB RESOURCES

APSIS (Analysis of Power for Sequencing-and-Imputation Studies): http://github.com/corbinq/APSIS

## ACKNOWLEDGMENTS

The authors acknowledge support from NIH grant R01 HG009976 (MB), the Austrian Science Fund (FWF) grant J-3401 (CF), NIH contracts N01-AG-1-2109 and HHSN271201100005C (FC), and The Fondazione di Sardegna, Prot. U1301.2015/ AI.1157.BE Prat. 2015-1651 (FC).

## TOPMed

Whole genome sequencing (WGS) for the Trans-Omics in Precision Medicine (TOPMed) program was supported by the National Heart, Lung and Blood Institute (NHLBI). WGS for NHLBI TOPMed: Genes-Environments and Admixture and Latino Americans (GALA II) Study (phs000920.v1.p1) was performed at the New York Genome Center (3R01HL117004-01S3). WGS for NHLBI TOPMed: The Jackson Heart Study (phs000964.v1.p1) was performed at the University of Washington Northwest Genomics Center (HHSN268201100037C). WGS for NHLBI TOPMed: The Genetic Epidemiology of Asthma in Costa Rica (phs000988.v1.p1) was performed at the University of Washington Northwest Genomics Center (3R37HL066289-13S1). Centralized read mapping and genotype calling, along with variant quality metrics and filtering were provided by the TOPMed Informatics Research Center (3R01HL-117626-02S1). Phenotype harmonization, data management, sample-identity QC, and general study coordination, were provided by the TOPMed Data Coordinating Center (3R01HL-120393-02S1). We gratefully acknowledge the studies and participants who provided biological samples and data for TOPMed.

### Jackson Heart Study

The Jackson Heart Study (JHS) is supported and conducted in collaboration with Jackson State University (HHSN268201300049C and HHSN268201300050C), Tougaloo College (HHSN268201300048C), and the University of Mississippi Medical Center (HHSN268201300046C and HHSN268201300047C) contracts from the National Heart, Lung, and Blood Institute (NHLBI) and the National Institute for Minority Health and Health Disparities (NIMHD). The authors also wish to thank the staffs and participants of the JHS.

### The Genetics of Asthma in Costa Rica Project

The Genetics of Asthma in Costa Rica project is supported by R-37 HL066289 (PI-Scott T. Weiss) and P01 HL132825-01 (PI-Scott T. Weiss) both from the National Heart Lung and Blood Institute. The investigators wish to acknowledge the families who contributed data to this study and to Juan C.Celedon, MD, DPH, Lydia Avila MD, and Manuel Soto-Queros MD who led the data collection team in Costa Rica.

### Genes-Environments and Admixture in Latino Americans Study

The Genes-Environments and Admixture in Latino Americans (GALA II) Study and E.G.B. are supported by the Sandler Family Foundation, the American Asthma Foundation, the RWJF Amos Medical Faculty Development Program, the Harry Wm. and Diana V. Hind Distinguished Professor in Pharmaceutical Sciences II, the National Heart, Lung, and Blood Institute (NHLBI) R01HL117004, R01HL128439, R01HL135156, X01HL134589, the National Institute of Environmental Health Sciences R01ES015794, R21ES24844, the National Institute on Minority Health and Health Disparities (NIMHD) P60MD006902, R01MD010443, RL5GM118984 and the Tobacco-Related Disease Research Program under Award Number 24RT-0025. The authors wish to acknowledge the following GALA II co-investigators for subject recruitment, sample processing and quality control: Celeste Eng, Scott Huntsman, MSc, Donglei Hu, PhD, Angel CY Mak, MPhil, PhD, Shannon Thyne, MD, Harold J. Farber, MD, MSPH, Pedro C. Avila, MD, Denise Serebrisky, MD, William Rodriguez-Cintron, MD, Jose R. Rodriguez-Santana, MD, Rajesh Kumar, MD, Luisa N. Borrell, DDS, PhD, Emerita Brigino-Buenaventura, MD, Adam Davis, MA, MPH, Michael A. LeNoir, MD, Kelley Meade, MD, Saunak Sen, PhD and Fred Lurmann, MS. The authors also wish to thank the staffs and participants contributed to the GALA II study.

### SISu and Kuusamo

The SISu and Kuusamo dataset comprise of selected study samples from the FINRISK and Health2000 (H2000) studies. FINRISK study has been primarily funded by budgetary funds of THL (National Institute for Health and Welfare) with important additional funding from the Academy of Finland (grant number 139635 for VS) and from the Finnish Foundation for Cardiovascular Research. The H2000 study was funded by the National Institute for Health and Welfare (THL), the Finnish Centre for Pensions (ETK), the Social Insurance Institution of Finland (KELA), the Local Government Pensions Institution (KEVA) and other organizations listed on the website of the survey (http://www.terveys2000.fi). Both studies are grateful for the THL DNA laboratory for its skillful work to produce the DNA samples used in this study. We thank the Sanger Institute sequencing facilities for whole genome sequencing of the Kuusamo subset. AP and SR are supported by the Academy of Finland (grant no. 251704, 286500, 293404 to AP, and 251217, 285380 to SR), Juselius Foundation, Finnish Foundation for Cardiovascular Research, NordForsk eScience NIASC (grant no 62721) and Biocentrum Helsinki (to SR).

### GoT2D

Funding for this study was provided by: The Academy of Finland (139635); Action on Hearing Loss (UK) (G51); The Ahokas Foundation; The Andrea and Charles Bronfman Philanthropies; The British Heart Foundation (SP/04/002); The Canadian Institutes of Health Research; The Estonian Government (IUT24-6, IUT20-60); The European Commission (ENGAGE: HEALTH-F4-2007-201413); The European Community’s Seventh Framework Programme (FP7/2007-2013) (EpiMigrant, 279143); The European Regional Development Fund (3.2.0304.11-0312); The European Research Council (ADG20110310#293574); The European Union (EXGENESIS); The Finnish Diabetes Research Foundation; The Finnish Foundation for Cardiovascular Research; The Finnish Medical Society; The Folkhälsan Research Foundation (Finland); The Fonds de la Recherche en Santé du Québec (Canada); The Foundation for Life and Health in Finland; The German Federal Ministry of Education and Research (BMBF) The German Federal Ministry of Health (BMG); The German Center for Diabetes Research (DZD); Helmholtz Zentrum München (German Research Center for Environmental Health), which is supported by the German Federal Ministry of Education and Research (BMBF) and by the State of Bavaria; The Helsinki University Central Hospital Research Foundation; The KG Jebsen Foundation (Norway); The Knut och Alice Wallenberg Foundation (Sweden); The Medical Research Council (UK) (G0601261, G0601966, G0700931, G0900747); The Ministry of Science and Research of the State of North Rhine-Westphalia (MIWF NRW); The Munich Center of Health Sciences (MC-Health); The Närpes Health Care Foundation (Finland); The National Institute for Health Research (UK) (RP-PG-0407-10371); The National Institutes of Health (USA) (DK020595, DK062370, DK072193, DK073541, DK080140, DK085501, DK085526, DK085545, DK088389, DK092251, DK093757, DK098032, GM007753, HG005773, HHSN268201300046C, HHSN268201300047C, HHSN268201300048C, HHSN268201300049C, HHSN268201300050C, HL102830, MH090937, MH101820); The Nordic Center of Excellence in Disease Genetics; Novo Nordisk; The Ollqvist Foundation (Finland); The Paavo Nurmi Foundation (Finland); The Påhlssons Foundation (Sweden); The Perklén Foundation (Finland); The Research Council of Norway; The Signe and Ane Gyllenberg Foundation (Finland); The Sigrid Juselius Foundation (Finland); The Skåne Regional Health Authority (Sweden); The Swedish Cultural Foundation in Finland; The Swedish Diabetes Foundation (2012-1397, 2013-024); The Swedish Heart-Lung Foundation (20140422); The Swedish Research Council; The University of Bergen; The University of Tartu (SP1GVARENG); Uppsala University; The Wellcome Trust (UK) (064890, 083948, 084723, 085475, 086596, 090367, 090532, 092447, 095101, 095552, 098017, 098051, 098381);The Western Norway Regional Health Authority (Helse Vest).

### Disclaimer

The views expressed in this manuscript are those of the authors and do not necessarily represent the views of the National Heart, Lung, and Blood Institute; the National Institutes of Health; or the U.S. Department of Health and Human Services.

