## Supplemental Materials for "Sequencing and Imputation in GWAS: Cost-Effective Strategies to Increase Power and Genomic Coverage Across Diverse Populations"

**Supplemental Figure 1. Imputation Coverage and $r^2$ as Functions of Population-Specific Reference Panel Size**

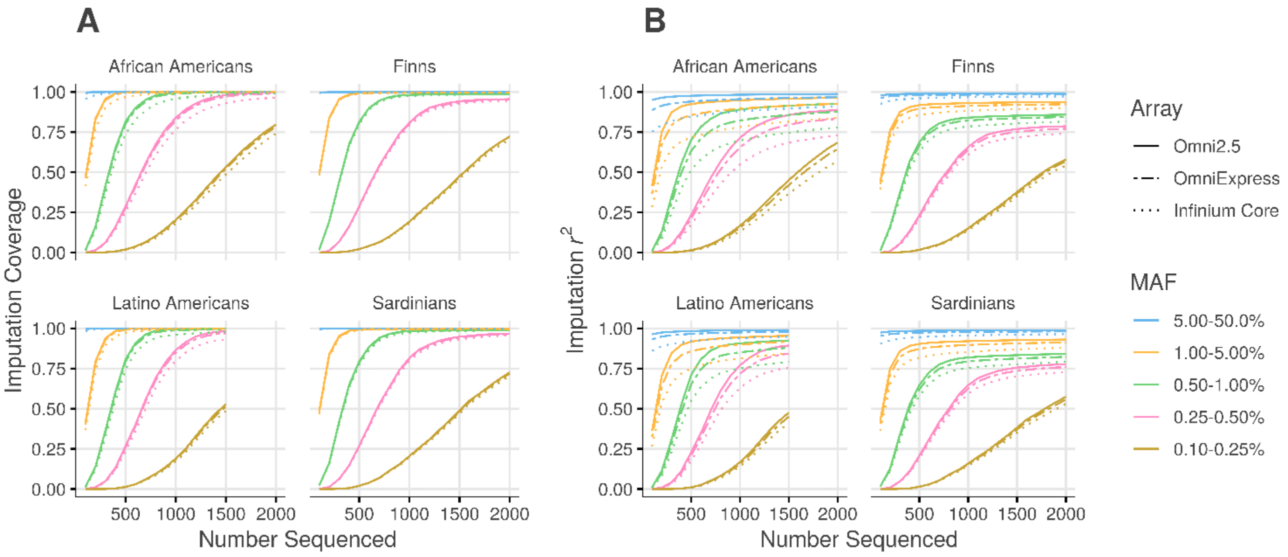

(**A**) Imputation coverage, defined as the proportion of variants with imputation $r^2 \geq 0.3$ and minor allele count (MAC) $\geq 5$ in the reference panel, and (**B**) imputation $r^2$, defined as the squared Pearson correlation between true genotype and imputed dosage, as a function of study-specific reference panel size (Number Sequenced).

**Supplemental Figure 2. Imputation $r^2$ for Augmented versus Distributed Reference Panels**

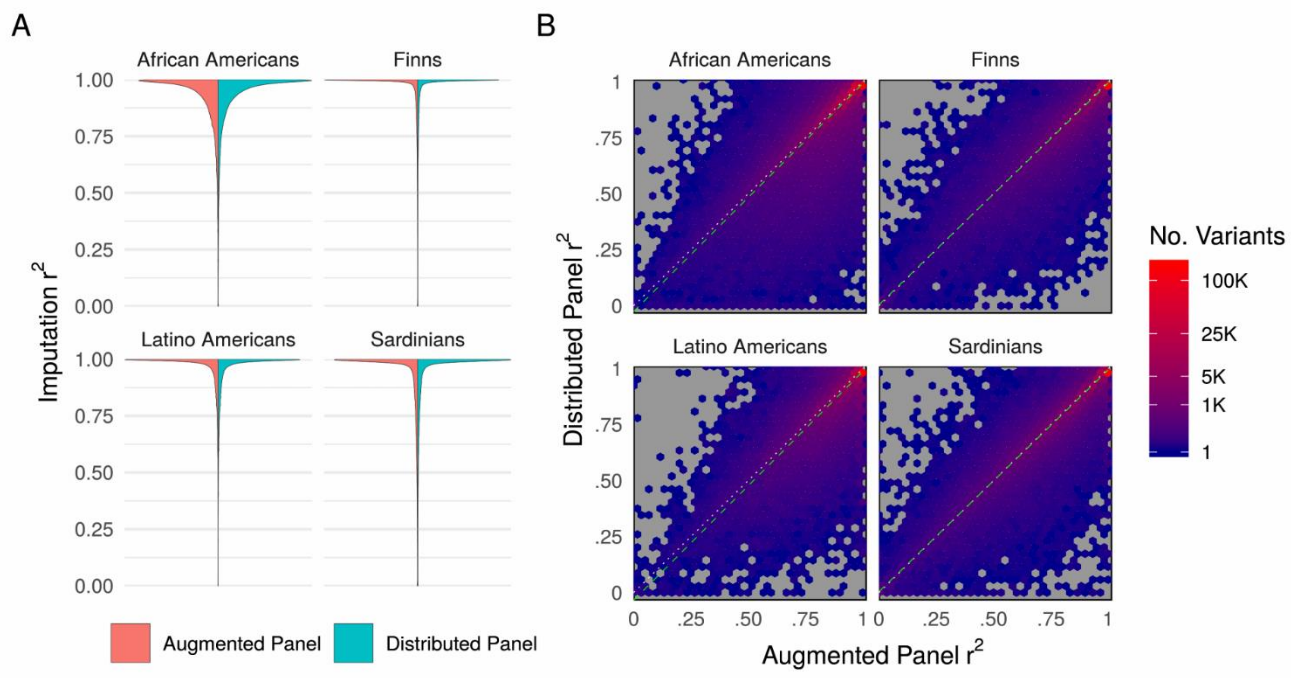

Imputation $r^2$ for augmented reference panels (directly augmenting the HRC or HRC subset with study-specific sequence data) versus a distributed reference panel approach, in which participants are separately imputed with the HRC (or HRC subset) and the study-specific reference panel and imputed variants from each panel are merged by selecting the set of imputed dosages with highest imputation quality (MaCH- $\hat{r}^2$) for each variant. Results are shown for the Omni Express array, and study specific reference panel sizes of 2,000 for African Americans, Finns, and Sardinians and 1,500 for Latino Americans. The marginal density of imputation $r^2$ for each population is shown in panel A, and binned scatterplots of imputation $r^2$ from augmented versus distributed panels is shown in panel B, with LSE regression lines in dashed green. The mean pairwise difference in imputation $r^2$ across variants between augmented and distributed panels was < 0.005 in magnitude for each population.

**Supplemental Figure 3. Optimal Designs for a Common Variant with Moderate Effect**

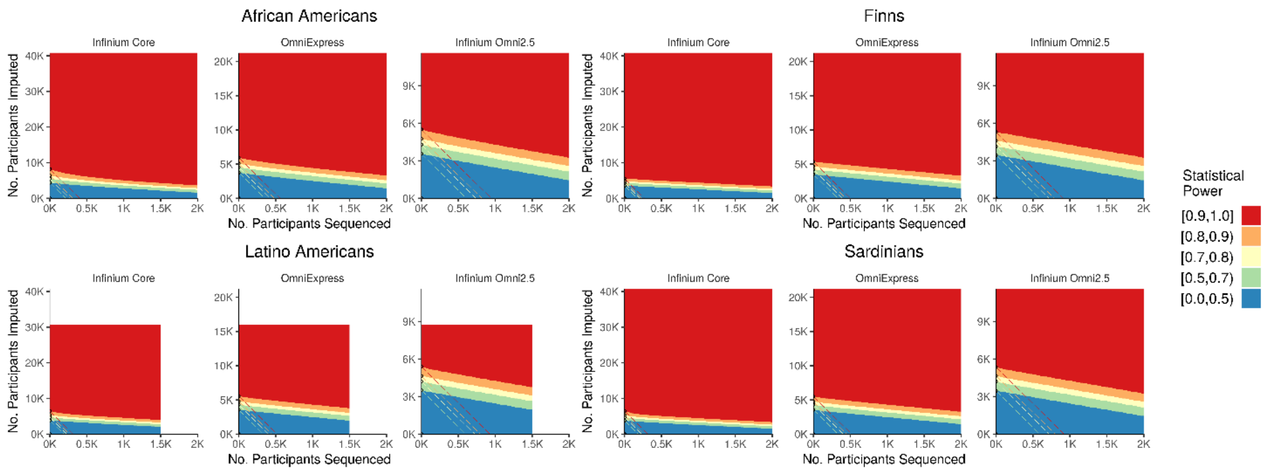

Power to detect association for case-control studies with equal numbers of cases and controls as a function of sequenced subsample size (x-axis) and imputed subsample size (y-axis) for a variant with MAF 10% and relative risk 1.5 for a disease with prevalence 1%. Axes scaled to reflect costs of genotyping arrays (Table 1) and sequencing ($1K per sample). Optimal designs are indicated only for total genotyping cost ≤ $2M ($1.5M for Latino Americans). Dashed diagonal lines indicate study designs with the same total cost, given by y=a-bx where a=(Total Cost) /(Array Cost) and b=(Sequencing Cost)/(Array Cost). Circled points indicate optimal study designs, which attain the indicated power level at minimum total experimental cost (or, maximize power at the indicated total experimental cost). In this example, exclusively array-based genotyping and imputing from the external HRC panel is optimal for all populations considered.

**Supplemental Figure 4. Optimal Designs for a Rare Variant with Large Effect**

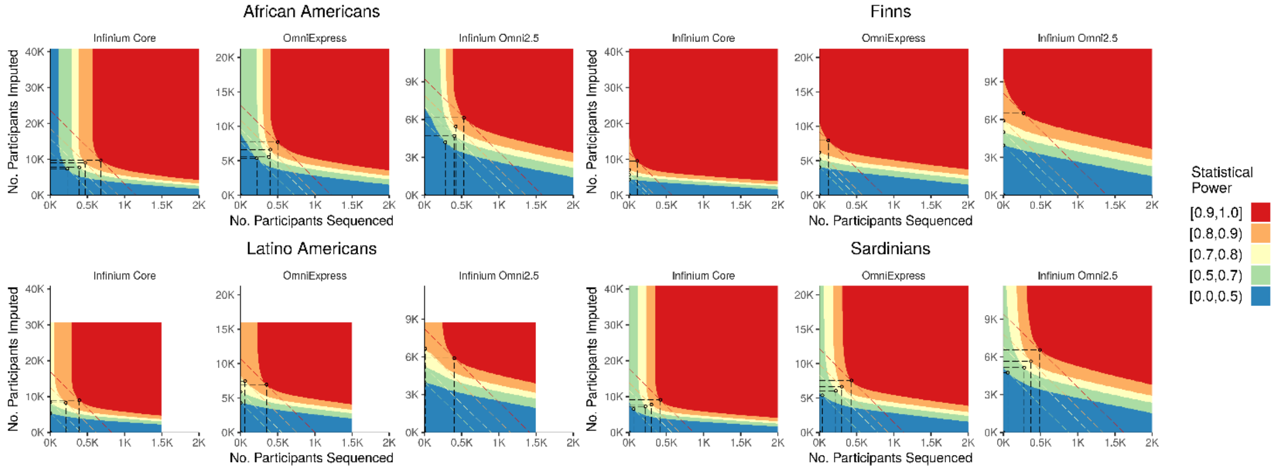

Power to detect association for case-control studies with equal numbers of cases and controls as a function of sequenced subsample size (x-axis) and imputed subsample size (y-axis) for a variant with MAF 0.25% and relative risk 6 for a disease with prevalence 1%. Axes scaled to reflect costs of genotyping arrays (Table 1) and sequencing ($1K per sample). Optimal designs are indicated only for total genotyping cost ≤ $2M ($1.5M for Latino Americans). Dashed diagonal lines indicate study designs with the same total cost, given by y=a-bx where a=(Total Cost) /(Array Cost) and b=(Sequencing Cost)/(Array Cost). Circled points indicate optimal study designs, which attain the indicated power level at minimum total experimental cost (or, maximize power at the indicated total experimental cost). In this example, sequencing a subset of participants is almost uniformly optimal across all populations.

### NHLBI Trans-Omics for Precision Medicine (TOPMed) Consortium Investigator List

| Name | Institution(s) |
|---|---|
| Abe, Namiko | New York Genome Center |
| Abecasis, Goncalo | University of Michigan |
| Albert, Christine | Massachusetts General Hospital |
| Allred, Nicholette (Nichole) Palmer | Wake Forest Baptist Health |
| Almasy, Laura | Children's Hospital of Philadelphia, University of Pennsylvania |
| Alonso, Alvaro | Emory University |
| Ament, Seth | University of Maryland |
| Anderson, Peter | University of Washington |
| Anugu, Pramod | University of Mississippi |
| Applebaum-Bowden, Deborah | National Institutes of Health |
| Arking, Dan | Johns Hopkins University |
| Arnett, Donna K | University of Kentucky |
| Ashley-Koch, Allison | Duke University |
| Aslibekyan, Stella | University of Alabama |
| Assimes, Tim | Stanford University |
| Auer, Paul | University of Wisconsin Milwaukee |
| Avramopoulos, Dimitrios | Johns Hopkins University |
| Barnard, John | Cleveland Clinic |
| Barnes, Kathleen | University of Colorado at Denver |
| Barr, R. Graham | Columbia University |
| Barron-Casella, Emily | Johns Hopkins University |

| Name | Institution(s) |
| --- | --- |
| Beaty, Terri | Johns Hopkins University |
| Becker, Diane | Johns Hopkins University |
| Becker, Lewis | Johns Hopkins University |
| Beer, Rebecca | National Institutes of Health |
| Begum, Ferdouse | Johns Hopkins University |
| Beitelshees, Amber | University of Maryland |
| Benjamin, Emelia | Boston University, Massachusetts General Hospital |
| Bezerra, Marcos | Fundação de Hematologia e Hemoterapia de Pernambuco - Hemope |
| Bielak, Larry | University of Michigan |
| Bis, Joshua | University of Washington |
| Blackwell, Thomas | University of Michigan |
| Blangero, John | University of Texas Rio Grande Valley School of Medicine |
| Boerwinkle, Eric | University of Texas Health |
| Borecki, Ingrid | University of Washington |
| Bowler, Russell | National Jewish Health |
| Brody, Jennifer | University of Washington |
| Broeckel, Ulrich | Medical College of Wisconsin |
| Broome, Jai | University of Washington |
| Bunting, Karen | New York Genome Center |
| Burchard, Esteban | University of California, San Francisco |
| Cardwell, Jonathan | University of Colorado at Denver |
| Carty, Cara | Women's Health Initiative |
| Casaburi, Richard | University of California, Los Angeles |

| Name | Institution(s) |
| --- | --- |
| Casella, James | Johns Hopkins University |
| Chaffin, Mark | The Broad Institute |
| Chang, Christy | University of Maryland |
| Chasman, Daniel | Brigham & Women's Hospital |
| Chavan, Sameer | University of Colorado at Denver |
| Chen, Bo-Juen | New York Genome Center |
| Chen, Wei-Min | University of Virginia |
| Chen, Yii-Der Ida | Los Angeles Biomedical Research Institute |
| Cho, Michael | Brigham & Women's Hospital |
| Choi, Seung Hoan | The Broad Institute |
| Chuang, Lee-Ming | National Taiwan University |
| Chung , Mina | Cleveland Clinic |
| Cornell, Elaine | University of Vermont |
| Correa, Adolfo | University of Mississippi |
| Crandall, Carolyn | University of California, Los Angeles |
| Crapo, James | National Jewish Health |
| Cupples, L Adrienne | Boston University |
| Curran, Joanne | University of Texas Rio Grande Valley School of Medicine |
| Curtis, Jeffrey | University of Michigan |
| Custer, Brian | Blood Systems Research Institute UCSF |
| Damcott, Coleen | University of Maryland |
| Darbar, Dawood | University of Illinois at Chicago |
| Das, Sayantan | University of Michigan |

| Name | Institution(s) |
| --- | --- |
| David, Sean | Stanford University |
| Davis, Colleen | University of Washington |
| Daya, Michelle | University of Colorado at Denver |
| de Andrade, Mariza | Mayo Clinic |
| DeBaun, Michael | Vanderbilt University |
| Deka, Ranjan | University of Cincinnati |
| DeMeo, Dawn | Brigham & Women's Hospital |
| Devine, Scott | University of Maryland |
| Do, Ron | Icahn School of Medicine at Mount Sinai |
| Duan, Qing | University of North Carolina |
| Duggirala, Ravi | University of Texas Rio Grande Valley School of Medicine |
| Durda, Peter | University of Vermont |
| Dutcher, Susan | Washington University in St Louis |
| Eaton, Charles | Brown University |
| Ekunwe, Lynette | University of Mississippi |
| Ellinor, Patrick | Massachusetts General Hospital |
| Emery, Leslie | University of Washington |
| Farber, Charles | University of Virginia |
| Farnam, Leanna | Brigham & Women's Hospital |
| Fingerlin, Tasha | National Jewish Health |
| Flickinger, Matthew | University of Michigan |
| Fornage, Myriam | University of Texas Health |
| Franceschini, Nora | University of North Carolina |

| Name | Institution(s) |
|---|---|
| Fu, Mao | University of Maryland |
| Fullerton, Malia | University of Washington |
| Fulton, Lucinda | Washington University in St Louis |
| Gabriel, Stacey | The Broad Institute |
| Gan, Weiniu | National Institutes of Health |
| Gao, Yan | University of Mississippi |
| Gass, Margery | Fred Hutchinson Cancer Research Center |
| Gelb, Bruce | Icahn School of Medicine at Mount Sinai |
| Geng, Xiaoqi (Priscilla) | University of Michigan |
| Germer, Soren | New York Genome Center |
| Gignoux, Chris | Stanford University |
| Gladwin, Mark | University of Pittsburgh |
| Glahn, David | Yale University |
| Gogarten, Stephanie | University of Washington |
| Gong, Da-Wei | University of Maryland |
| Goring, Harald | University of Texas Rio Grande Valley School of Medicine |
| Gu, C. Charles | Washington University in St Louis |
| Guan, Yue | University of Maryland |
| Guo, Xiuqing | Los Angeles Biomedical Research Institute |
| Haessler, Jeff | Fred Hutchinson Cancer Research Center, Women's Health Initiative |
| Hall, Michael | University of Mississippi |
| Harris, Daniel | University of Maryland |
| Hawley, Nicola | Yale University |

| Name | Institution(s) |
| --- | --- |
| He, Jiang | Tulane University |
| Heavner, Ben | University of Washington |
| Heckbert, Susan | University of Washington |
| Hernandez, Ryan | University of California, San Francisco |
| Herrington, David | Wake Forest Baptist Health |
| Hersh, Craig | Brigham & Women's Hospital |
| Hidalgo, Bertha | University of Alabama |
| Hixson, James | University of Texas Health |
| Hokanson, John | University of Colorado at Denver |
| Hong, Elliott | University of Maryland |
| Hoth, Karin | University of Iowa |
| Hsiung, Chao (Agnes) | National Health Research Institute Taiwan |
| Huston, Haley | Blood Works Northwest |
| Hwu, Chii Min | Taichung Veterans General Hospital Taiwan |
| Irvin, Marguerite Ryan | University of Alabama |
| Jackson, Rebecca | Ohio State University Wexner Medical Center |
| Jain, Deepti | University of Washington |
| Jaquish, Cashell | National Institutes of Health |
| Jhun, Min A | University of Michigan |
| Johnsen, Jill | Blood Works Northwest, University of Washington |
| Johnson, Andrew | NIH National Heart, Lung, and Blood Institute |
| Johnson, Craig | University of Washington |
| Johnston, Rich | Emory University |

| Name | Institution(s) |
| --- | --- |
| Jones, Kimberly | Johns Hopkins University |
| Kang, Hyun Min | University of Michigan |
| Kaplan, Robert | Albert Einstein College of Medicine |
| Kardia, Sharon | University of Michigan |
| Kathiresan, Sekar | The Broad Institute |
| Kaufman, Laura | Brigham & Women's Hospital |
| Kelly, Shannon | Blood Systems Research Institute UCSF |
| Kenny, Eimear | Icahn School of Medicine at Mount Sinai |
| Kessler, Michael | University of Maryland |
| Khan, Alyna | University of Washington |
| Kinney, Greg | University of Colorado at Denver |
| Konkle, Barbara | Blood Works Northwest |
| Kooperberg, Charles | Fred Hutchinson Cancer Research Center |
| Kramer, Holly | Loyola University |
| Krauter, Stephanie | University of Washington |
| Lange, Christoph | Harvard School of Public Health |
| Lange, Ethan | University of Colorado at Denver |
| Lange, Leslie | University of Colorado at Denver |
| Laurie, Cathy | University of Washington |
| Laurie, Cecelia | University of Washington |
| LeBoff, Meryl | Brigham & Women's Hospital |
| Lee, Seunggeun Shawn | University of Michigan |
| Lee, Wen-Jane | Taichung Veterans General Hospital Taiwan |

| Name | Institution(s) |
| --- | --- |
| LeFaive, Jonathon | University of Michigan |
| Levine, David | University of Washington |
| Levy, Dan | NIH National Heart, Lung, and Blood Institute, National Institutes of Health |
| Lewis, Joshua | University of Maryland |
| Li, Yun | University of North Carolina |
| Lin, Honghuang | Boston University |
| Lin, Keng Han | University of Michigan |
| Liu, Simin | Brown University, Women's Health Initiative |
| Liu, Yongmei | Wake Forest Baptist Health |
| Loos, Ruth | Icahn School of Medicine at Mount Sinai |
| Lubitz, Steven | Massachusetts General Hospital |
| Lunetta, Kathryn | Boston University |
| Luo, James | NIH National Heart, Lung, and Blood Institute, National Institutes of Health |
| Mahaney, Michael | University of Texas Rio Grande Valley School of Medicine |
| Make, Barry | Johns Hopkins University |
| Manichaikul, Ani | University of Virginia |
| Manson, JoAnn | Brigham & Women's Hospital |
| Margolin, Lauren | The Broad Institute |
| Martin, Lisa | George Washington University |
| Mathai, Susan | University of Colorado at Denver |
| Mathias, Rasika | Johns Hopkins University |
| McArdle, Patrick | University of Maryland |

| Name | Institution(s) |
|---|---|
| McDonald, Merry-Lynn | University of Alabama |
| McFarland, Sean | Harvard University |
| McGarvey, Stephen | Brown University |
| Mei, Hao | University of Mississippi |
| Meyers, Deborah A | University of Arizona |
| Mikulla, Julie | National Institutes of Health |
| Min, Nancy | University of Mississippi |
| Minear, Mollie | National Institutes of Health |
| Minster, Ryan L | University of Pittsburgh |
| Mitchell, Braxton | University of Maryland |
| Montasser, May E. | University of Maryland |
| Musani, Solomon | University of Mississippi |
| Mwasongwe, Stanford | University of Mississippi |
| Mychaleckyj, Josyf C | University of Virginia |
| Nadkarni, Girish | Icahn School of Medicine at Mount Sinai |
| Naik, Rakhi | Johns Hopkins University |
| Natarajan, Pradeep | The Broad Institute, Harvard University, Massachusetts General Hospital |
| Nekhai, Sergei | Howard University |
| Nickerson, Deborah | University of Washington |
| North, Kari | University of North Carolina |
| O'Connell, Jeff | University of Maryland |
| O'Connor, Tim | University of Maryland |
| Ochs-Balcom, Heather | University at Buffalo |

| Name | Institution(s) |
| --- | --- |
| Pankow, James | University of Minnesota |
| Papanicolaou, George | National Institutes of Health |
| Parker, Margaret | Brigham & Women's Hospital |
| Parsa, Afshin | University of Maryland |
| Penchev, Sara | National Jewish Health |
| Peralta, Juan Manuel | University of Texas Rio Grande Valley School of Medicine |
| Perez, Marco | Stanford University |
| Perry, James | University of Maryland |
| Peters, Ulrike | Fred Hutchinson Cancer Research Center, University of Washington |
| Peyser, Patricia | University of Michigan |
| Phillips, Larry | Emory University |
| Phillips, Sam | University of Washington |
| Pollin, Toni | University of Maryland |
| Post, Wendy | Johns Hopkins University |
| Powers Becker, Julia | University of Colorado at Denver |
| Preethi Boorgula, Meher | University of Colorado at Denver |
| Preuss, Michael | Icahn School of Medicine at Mount Sinai |
| Prokopenko, Dmitry | Harvard University |
| Psaty, Bruce | University of Washington |
| Qasba, Pankaj | National Institutes of Health |
| Qiao, Dandi | Brigham & Women's Hospital |
| Qin, Zhaohui | Emory University |
| Rafaels, Nicholas | University of Colorado at Denver |

| Name | Institution(s) |
| --- | --- |
| Raffield, Laura | University of North Carolina |
| Ramachandran, Vasan | Boston University |
| Rao, D.C. | Washington University in St Louis |
| Rasmussen-Torvik, Laura | Northwestern University |
| Ratan, Aakrosh | University of Virginia |
| Redline, Susan | Brigham & Women's Hospital |
| Reed, Robert | University of Maryland |
| Regan, Elizabeth | National Jewish Health |
| Reiner, Alex | Fred Hutchinson Cancer Research Center, University of Washington |
| Rice, Ken | University of Washington |
| Rich, Stephen | University of Virginia |
| Roden, Dan | Vanderbilt University |
| Roselli, Carolina | The Broad Institute |
| Rotter, Jerome | Los Angeles Biomedical Research Institute |
| Ruczinski, Ingo | Johns Hopkins University |
| Russell, Pamela | University of Colorado at Denver |
| Ruuska, Sarah | Blood Works Northwest |
| Ryan, Kathleen | University of Maryland |
| Sakornsakolpat, Phuwanat | Brigham & Women's Hospital |
| Salimi, Shabnam | University of Maryland |
| Salzberg, Steven | Johns Hopkins University |
| Sandow, Kevin | Los Angeles Biomedical Research Institute |
| Sankaran, Vijay | Harvard University |

| Name | Institution(s) |
|---|---|
| Scheller, Christopher | University of Michigan |
| Schmidt, Ellen | University of Michigan |
| Schwander, Karen | Washington University in St Louis |
| Schwartz, David | University of Colorado at Denver |
| Sciurba, Frank | University of Pittsburgh |
| Seidman, Christine | Harvard Medical School |
| Sheehan, Vivien | Baylor College of Medicine |
| Shetty, Amol | University of Maryland |
| Shetty, Aniket | University of Colorado at Denver |
| Sheu, Wayne Hui-Heng | Taichung Veterans General Hospital Taiwan |
| Shoemaker, M. Benjamin | Vanderbilt University |
| Silver, Brian | University of Massachusetts Memorial Health Center |
| Silverman, Edwin | Brigham & Women's Hospital |
| Smith, Jennifer | University of Michigan |
| Smith, Josh | University of Washington |
| Smith, Nicholas | University of Washington |
| Smith, Tanja | New York Genome Center |
| Smoller, Sylvia | Albert Einstein College of Medicine |
| Snively, Beverly | Wake Forest Baptist Health |
| Sofer, Tamar | Brigham & Women's Hospital |
| Sotoodehnia, Nona | University of Washington |
| Stilp, Adrienne | University of Washington |
| Streeten, Elizabeth | University of Maryland |

| Name | Institution(s) |
| --- | --- |
| Sung, Yun Ju | Washington University in St Louis |
| Sylvia, Jody | Brigham & Women's Hospital |
| Szpiro, Adam | University of Washington |
| Sztalryd, Carole | University of Maryland |
| Taliun, Daniel | University of Michigan |
| Tang, Hua | Stanford University |
| Taub, Margaret | Johns Hopkins University |
| Taylor, Kent | Los Angeles Biomedical Research Institute |
| Taylor, Simeon | University of Maryland |
| Telen, Marilyn | Duke University |
| Thornton, Timothy A. | University of Washington |
| Tinker, Lesley | Women's Health Initiative |
| Tirschwell, David | University of Washington |
| Tiwari, Hemant | University of Alabama |
| Tracy, Russell | University of Vermont |
| Tsai, Michael | University of Minnesota |
| Vaidya, Dhananjay | Johns Hopkins University |
| VandeHaar, Peter | University of Michigan |
| Vrieze, Scott | University of Colorado at Boulder, University of Minnesota |
| Walker, Tarik | University of Colorado at Denver |
| Wallace, Robert | University of Iowa |
| Walts, Avram | University of Colorado at Denver |
| Wan, Emily | Brigham & Women's Hospital |

| Name | Institution(s) |
|---|---|
| Wang, Fei Fei | University of Washington |
| Watson, Karol | University of California, Los Angeles |
| Weeks, Daniel E. | University of Pittsburgh |
| Weir, Bruce | University of Washington |
| Weiss, Scott | Brigham & Women's Hospital |
| Weng, Lu-Chen | Massachusetts General Hospital |
| Willer, Cristen | University of Michigan |
| Williams, Kayleen | University of Washington |
| Williams, L. Keoki | Henry Ford Health System |
| Wilson, Carla | Brigham & Women's Hospital |
| Wilson, James | University of Mississippi |
| Wong, Quenna | University of Washington |
| Xu, Huichun | University of Maryland |
| Yanek, Lisa | Johns Hopkins University |
| Yang, Ivana | University of Colorado at Denver |
| Yang, Rongze | University of Maryland |
| Zaghloul, Norann | University of Maryland |
| Zhang, Yingze | University of Pittsburgh |
| Zhao, Snow Xueyan | National Jewish Health |
| Zhao, Wei | University of Michigan |
| Zheng, Xiuwen | University of Washington |
| Zhi, Degui | University of Texas Health |
| Zhou, Xiang | University of Michigan |

| Name | Institution(s) |
| --- | --- |
| Zody, Michael | New York Genome Center |
| Zoellner, Sebastian | University of Michigan |
